# TRACE: Trajectory Reconstruction and Clustering from Sparse Evidence for Disease Subtyping and Staging

**DOI:** 10.64898/2026.11.11.765968

**Authors:** Yuanwang Zhang, Hongming Li, Cong Wang, Guoliang You, Yong Fan

**Affiliations:** Department of Bioengineering, University of Pennsylvania, Philadelphia, PA, USA; Department of Radiology, Perelman School of Medicine, University of Pennsylvania, Philadelphia, PA, USA

## Abstract

Chronic progressive diseases often follow heterogeneous courses, with individuals differing in the pattern and timing of biomarker abnormalities. However, observed data are typically sparse and predominantly cross-sectional, with limited follow-up covering only short segments of disease progression. Reconstructing heterogeneous trajectories from such data requires disentangling variation along disease trajectories from subtype heterogeneity while inferring individuals’ unknown sub-type assignments and disease stages. Existing approaches may confound disease stage with subtype, scale poorly as the number of biomarkers increases, or fail to effectively integrate cross-sectional and longitudinal observations. We propose ***TRACE*** (**T**rajectory **R**econstruction **A**nd **C**lustering from Sparse **E**vidence), a probabilistic framework that jointly reconstructs biomarker trajectories and infers disease subtypes and continuous stages. TRACE combines monotonic neural networks for flexible modeling and data-driven clustering of subtype-specific progression trajectories with temporal regularization derived from longitudinal observations when available. All model components are jointly optimized end to end using stochastic gradient descent, enabling efficient and scalable estimation. We evaluate TRACE on semi-simulated MRI biomarker data with known subtype and stage structures and demonstrate its ability to identify clinically meaningful subtypes and disease stages in real-world neurodegenerative disease datasets.

## 1 Introduction

Understanding chronic disease progression requires characterizing both how biomarker profiles evolve over time and how their trajectories differ across individuals. This is particularly challenging for diseases that progress over years or decades, as individuals with the same clinical diagnosis may follow distinct courses that differ in the biomarkers affected, the timing of abnormalities, and the rate of progression. Identifying these patterns can support patient stratification and inform prognosis (Ahlqvist et al., 2018). In this study, we focus on neurodegenerative diseases and use MRI biomarkers to characterize changes in brain structure. These diseases may progress over decades, whereas individual follow-up typically captures only a small portion of the disease course (Bateman et al., 2012; Young et al., 2024). Ideally, progression trajectories would be reconstructed from dense longitudinal observations spanning the entire disease course. In practice, however, available data are often sparse and predominantly cross-sectional, with only a small number of repeated measurements collected over short intervals (Donohue et al., 2014), as schematically illustrated in Fig. 1(a). Because individuals are observed at different and unknown disease stages, these data provide neither a common disease timeline nor complete individual trajectories, making it difficult to reconstruct long-term disease progression and identify distinct progression subtypes.

**Figure 1:**
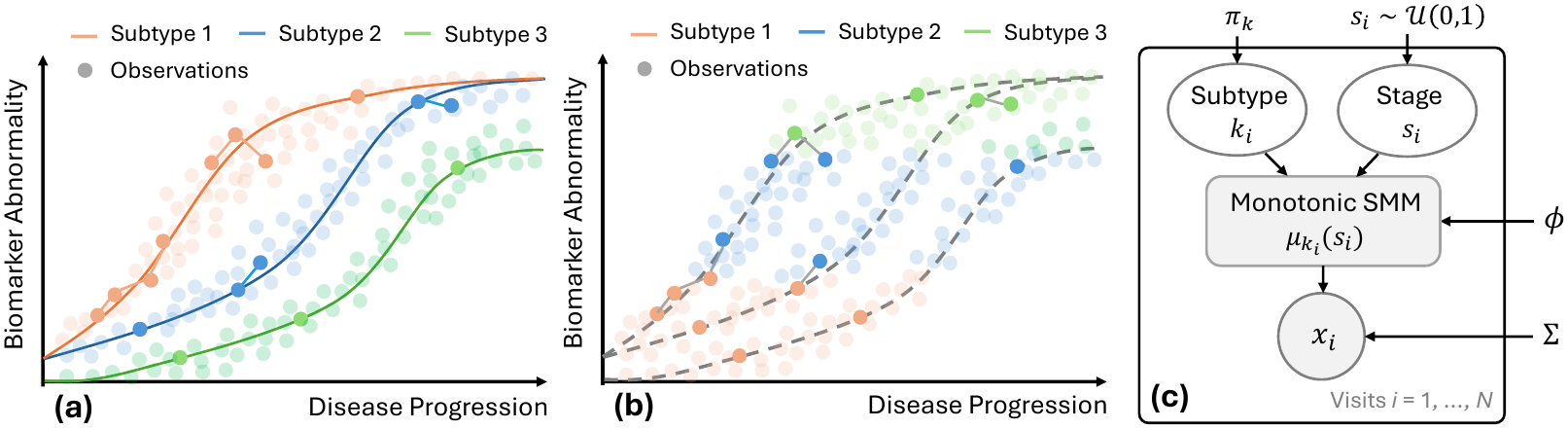
Motivation and overview of the proposed framework. (a) Distinct disease progression trajectories are observed through cross-sectional snapshots and short longitudinal segments. Multiple visits of the same individuals are connected by solid lines. (b) Conventional clustering may group participants by disease stage rather than progression subtype. (c) Graphical model of the proposed probabilistic subtype-and-stage framework.

Conventional clustering methods (Dong et al., 2015; Varol et al., 2017; Yang et al., 2021; Kang et al., 2024) do not adequately address this problem. When applied directly to cross-sectional measurements, these methods group individuals based on the similarity of their observed biomarker profiles without accounting for their position along the disease course, as schematically illustrated in Fig. 1(b). Consequently, individuals following the same disease trajectory but at early and late stages may be assigned to different clusters, whereas those following distinct disease trajectories may appear similar at certain stages. The resulting clusters may therefore reflect disease severity rather than distinct progression patterns. Jointly modeling subtypes and stages is essential for disentangling variation along a progression trajectory from heterogeneity across trajectories.

Several methods have been developed to address this challenge. Subtype and Stage Inference (SuS-taIn) and its variants jointly estimate progression subtypes and stages from cross-sectional data by representing each subtype as an ordered sequence of biomarker abnormalities (Young et al., 2018; 2021; Tandon et al., 2024; de Jonge et al., 2026). However, searching over possible event orderings becomes computationally demanding as the number of biomarkers and events increases, limiting its scalability to fine-grained brain regional measurements. Surreal-GAN learns continuous disease-related imaging dimensions and individual severity indices from reference and patient samples (Yang et al., 2022; 2024). By allowing multiple disease-related patterns to coexist within an individual, its continuous-dimensional representation differs from discrete subtype models and may be less directly suited to identifying discrete subtypes. Moreover, existing formulations of SuStaIn and Surreal-GAN rely primarily on cross-sectional data and do not explicitly leverage temporal information from short-term longitudinal observations, potentially limiting their ability to distinguish progression patterns. SubLign instead aligns and clusters interval-censored multivariate time series (Chen et al., 2022). However, it is designed for longitudinal timeseries and therefore does not address the common setting in which many individuals have only a single observation. DPMoSt jointly models disease progression and stratifies individuals using sigmoid-shaped biomarker trajectories and subject-specific time shifts, allowing individual biomarkers to follow shared or subtype-specific progression patterns (Viani et al., 2025). However, its trajectories are restricted to a parametric sigmoid family, which may limit its flexibility in modeling complex biomarker progression dynamics. Together, these limitations motivate an efficient, scalable, and flexible framework that jointly estimates progression subtypes and continuous stages from cross-sectional observations while incorporating temporal information from short-term longitudinal data when available.

In this study, we propose **T**rajectory **R**econstruction **A**nd **C**lustering from Sparse **E**vidence (***TRACE***), a probabilistic framework in which each disease subtype is represented by a multivariate biomarker trajectory over continuous disease stages. By marginalizing over latent subtype assignments and disease stages in the observation likelihood, TRACE can learn from cross-sectional observations for estimating individuals’ posterior subtype probabilities and continuous stages. To flexibly model potentially complex disease progression trajectories, we parameterize each subtype-specific trajectory using a smooth min–max (SMM) monotonic neural network (Igel, 2024). When repeated visits are available, a change reconstruction loss incorporates longitudinal information by encouraging the predicted magnitude and pattern of biomarker changes to match those observed between adjacent visits. All model components are jointly optimized end to end using stochastic gradient descent, enabling efficient and scalable estimation. We first evaluate TRACE on semi-simulated MRI data with known subtype and stage ground truth, comparing its recovery performance with existing methods under both cross-sectional and longitudinal settings. We then apply TRACE to real-world neurodegenerative disease MRI datasets to assess whether the inferred trajectories and subtypes exhibit different patterns and clinically meaningful associations. TRACE achieves the best overall subtype and stage recovery on the semi-simulated data and identifies four distinct anatomical progression patterns associated with differences in clinical impairment in the real-world cohorts.

Our contributions are fourfold: (i) a probabilistic formulation that jointly infers disease subtypes and continuous stages from cross-sectional biomarker observations; (ii) flexible monotonic neural networks that model complex subtype-specific trajectories of the biomarker progression dynamics; (iii) a change reconstruction loss that incorporates temporal information from short-term longitudinal observations when available; and (iv) an end-to-end optimization strategy that avoids combinatorial event-ordering searches and enables computationally efficient, scalable estimation.

## 2 Method

### 2.1 Problem formulation

Let *i* ∈{1, …, *N}* index observations, with each observation corresponding to a single study visit. For each observation, we measure an *F*-dimensional biomarker vector ***x***_*i*_ ∈ R^*F*^, where larger values indicate greater abnormality. We assume that each observation is associated with two latent random variables: a disease subtype k_*i*_ *∈ {*1, …, *K}* and a continuous disease stage s_*i*_ *∈* [0, 1].

Each subtype *k* is characterized by a multivariate trajectory

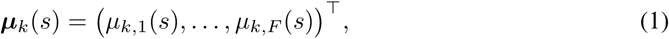

which represents the expected biomarker profile for subtype *k* at stage *s*. For each subtype *k* and biomarker *f*, we impose *μ*_*k,f*_ (0) = 0 and *∂μ*_*k,f*_ (*s*)*/∂s >* 0 such that trajectories begin at the reference-control mean and progress monotonically toward greater abnormality, as described in Sec. 2.4.

Given the observed datase 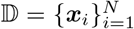, our goal is to learn the population-level subtype trajectories, subtype proportions, and observation noise. Given a learned model and an observed biomarker vector ***x***_*i*_, we infer its posterior subtype probabilities and latent disease-stage distribution, as described in Sec. 2.3. All observations contribute to the generative likelihood in Sec. 2.2. When repeated observations from the same individuals are available, they provide additional temporal supervision through the change reconstruction loss described in Sec. 2.5.

### 2.2 Observation model

We assign each subtype a prior probability

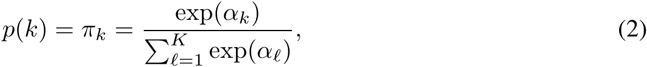

where 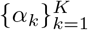 are learned logits. We place a uniform prior on the continuous disease stage, s_*i*_ *∼ U* (0, 1). Conditioned on subtype *k* and stage *s*, the biomarkers are modeled as conditionally independent Gaussian observations:

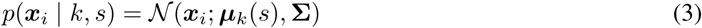

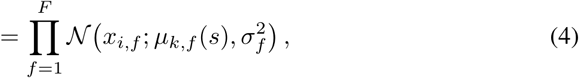

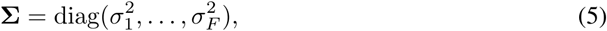

where each feature-specific noise scale *σ*_*f*_ is learned and shared across subtypes. Marginalizing over both latent variables yields the observation likelihood:

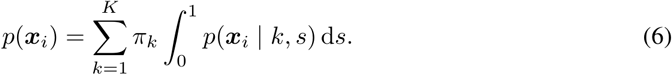

The observation loss is defined as the mean negative log-likelihood:

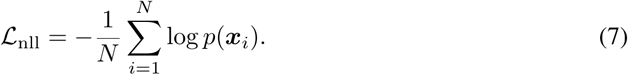

In practice, we approximate the stage integral using Gauss–Legendre quadrature. Given quadrature nodes 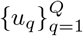 and weights 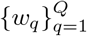 on [0, 1], Eq. 6 becomes

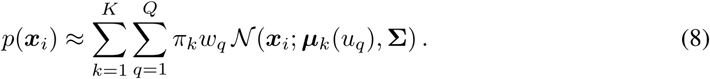

We compute the resulting log-likelihood using the log-sum-exp for numerical stability and use the same quadrature approximation for posterior inference.

### 2.3 Posterior inference of subtype and stage

Given the biomarker vector ***x***_*i*_ from an individual visit, Bayes’ rule yields a normalized joint posterior mass over subtype *k* and quadrature stage *u*_*q*_:

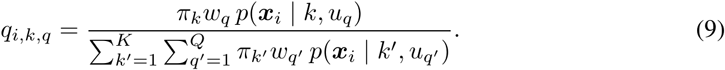

Marginalizing the joint mass yields the subtype posterior probabilities and the posterior mass assigned to each stage node:

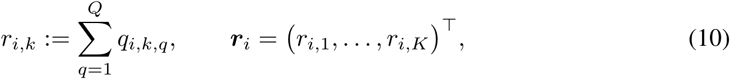

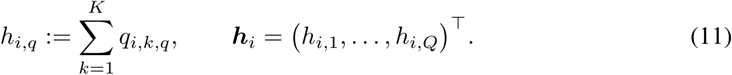

Thus, *r*_*i,k*_ approximates *P* (k_*i*_ = *k* |**x**_*i*_), whereas *h*_*i,q*_ approximates the posterior mass associated with stage node *u*_*q*_. We assign each observation to its maximum posterior subtype and estimate its disease stage using the posterior mean:

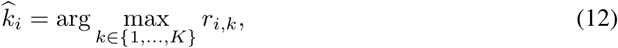

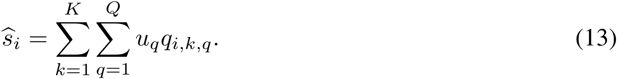

### 2.4 Monotonic neural networks for subtype-specific trajectories

Following previous work (Young et al., 2018; Yang et al., 2024), we interpret disease stage as accumulated disease-related abnormality. After aligning biomarker directions, we therefore require the expected abnormality of each biomarker to increase monotonically with disease stage. This assumption establishes a consistent direction of progression and prevents trajectories from returning to previously visited biomarker states, which could make stage ambiguous when learning from cross-sectional or sparsely sampled longitudinal observations.

We model these trajectories using smooth min–max (SMM) neural networks (Igel, 2024). For each subtype *k*, the network takes a scalar disease stage *s* ∈ [0, 1] as input and outputs an *F*-dimensional vector of expected normalized biomarker values. Each subtype–biomarker pair has its own learnable parameters, allowing trajectories to differ in both shape and magnitude across subtypes and biomarkers. SMM networks combine positive-slope affine functions through smooth minimum and maximum operations that preserve monotonicity. Consequently, each expected biomarker values from the output increases monotonically with disease stage while retaining the flexibility to represent nonlinear progression patterns.

Let **g**_*k*_(*s*) ∈ R^*F*^ denote the network output for subtype *k*. We anchor each trajectory at the reference-control mean by subtracting its value at stage zero:

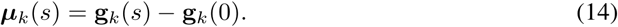

This construction ensures that ***μ***_*k*_(0) = **0** without altering monotonicity. The endpoint *μ*_*k,f*_ (1) is learned separately for every subtype and biomarker, allowing biomarkers with limited involvement in a given subtype to follow nearly flat trajectories.

We additionally implemented monotonic piecewise-linear and I-spline parameterizations as alternative trajectory models. Their definitions and corresponding experiments are provided in Appendix B.2.

### 2.5 Learning from short-term biomarker changes

The observation likelihood models each visit independently and therefore does not directly use biomarker changes observed across repeated visits from the same individual. We incorporate this temporal information through a change reconstruction loss that encourages the model to reproduce biomarker changes observed between adjacent visits.

Let *p* denote the set of chronologically ordered pairs (*i, j*) of adjacent visits from the same individual. Assuming that the two visits share a common disease subtype, we combine their visit-level subtype posteriors to obtain a shared pair-level distribution:

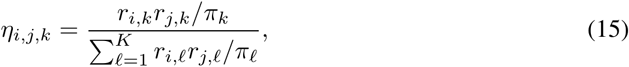

where *r*_*i,k*_ is the subtype posterior for visit *i*, calculated using Eq. 10. The resulting quantity *η*_*i,j,k*_represents the posterior support that the visit pair belongs to disease subtype *k*.

For each subtype, we reconstruct the expected biomarker profile by averaging its trajectory over the subtype-conditional stage posterior:

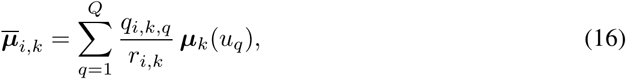

where *q*_*i,k,q*_ is the joint posterior mass defined in Eq. 9. For subtype *k*, the predicted change in biomarker *f* between visits *i* and *j* is 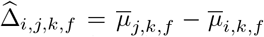, and the corresponding observed change is Δ_*i,j,f*_ = *x*_*j,f*_ *– x*_*i,f*_. We penalize their discrepancy using a posterior-weighted Huber loss:

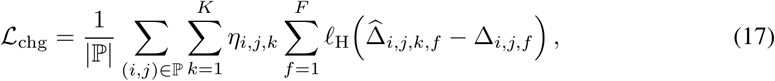

where *ℓ*_H_(*e*) = *e*^2^*/*2 for |*e*| ≤1 and *ℓ*_H_(*e*) = |*e*| −1*/*2 otherwise. The transition threshold is fixed at one normalized biomarker unit to reduce sensitivity to large residual changes. By comparing observed changes with differences between posterior-expected biomarker profiles, the loss provides temporal supervision without requiring known subtypes or stages.

The complete training objective is:

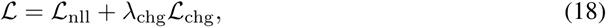

where *λ*_chg_ controls the contribution of the change reconstruction regularization. When no longitudinal visit pairs are available, the change-reconstruction term is omitted, allowing the model to be trained using cross-sectional observations alone.

### 2.6 Implementation details

#### Model parameters

The learned parameters include the subtype logits ***α***, the feature-specific noise parameters defining the diagonal covariance matrix σ, and the SMM weights and biases, collectively denoted by *ϕ*. All parameters are jointly optimized by minimizing Eq. 18, with gradients propagated through the likelihood, posterior computations, and change reconstruction loss.

#### Training details

Each SMM network contains four smooth-min groups with four affine units per group and uses a smoothing parameter of *β* = 10. We approximate the disease stage integral using eight-node Gauss–Legendre quadrature within each of seven equal subintervals of [0, 1], yielding 56 quadrature nodes in total. The models are implemented in PyTorch and trained using AdamW (Loshchilov & Hutter, 2019) with a learning rate of 3 ×10^−3^, a weight decay of 10^−5^, a mini-batch size of 256 observations, and gradient-norm clipping at 10. The subtype logits are initialized to zero, corresponding to uniform subtype prior probabilities. The SMM slopes are initialized near 0.5, and the biases are initialized using small zero-mean Gaussian perturbations.

## 3 Experiments and Results

### 3.1 Datasets and Preprocessing

#### Study cohorts

We used T1-weighted MRI data from the Alzheimer’s Disease Neuroimaging Initiative (ADNI) (Mueller et al., 2005) and the Open Access Series of Imaging Studies 3 (OASIS–3) (LaMontagne et al., 2019). We retained visits with complete information on age, sex, MRI biomarkers, and clinical diagnosis, including cognitively normal (CN), mild cognitive impairment (MCI), and dementia. The final dataset comprised 3,520 participants and 11,769 visits; additional cohort details are provided in Appendix A.1.

#### Image preprocessing

T1-weighted MRIs were processed using the standard FreeSurfer pipeline (Fischl, 2012) to extract 96 region-of-interest (ROI) biomarkers, including mean cortical thickness of 34 regions per hemisphere based on the Desikan–Killiany atlas (Desikan et al., 2006) and 28 volume measures obtained from FreeSurfer’s automated anatomical segmentation (aseg) (Fischl et al., 2002). Volume measures were normalized by total intracranial volume (TIV) to account for differences in overall head size.

#### Biomarker normalization

ROI measurements were transformed into *z*-scores so that features measured on different physical scales contribute comparably to likelihood and trajectory estimation. Normalization parameters were estimated from amyloid-negative CN individuals, who were less likely to harbor Alzheimer’s disease pathology than individuals at an early disease stage. To account for age- and sex-related variation in brain morphology, each raw feature was regressed on centered age and sex within the reference group. The resulting residuals were divided by their reference-group standard deviations. Feature directions were harmonized so that positive *z*-scores consistently indicate greater abnormality. Ventricular measures retained their original sign, whereas cortical thickness and other parenchymal measures were sign-reversed. Standardized values were clipped to [ −8, 8] to limit the influence of extreme observations. Reference participants were used to estimate normalization parameters but were excluded from subtype discovery.

### 3.2 Semi-simulated Data Experiments

Real neuroimaging datasets do not provide ground-truth disease subtypes, stages, or biomarker trajectories, precluding direct evaluation of their recovery. We therefore generated semi-simulated data that preserved the covariance and noise structure of real imaging measurements while providing known subtype assignments, disease stages, affected regions, and monotonic progression trajectories. This setting enabled controlled evaluation of both subtyping and staging recovery under the scientific assumptions used by the model.

#### Semi-simulated data generation

We constructed a semi-simulated dataset using baseline MRI measurements from 788 amyloid-negative cognitively normal participants in ADNI and OASIS–3. Participants were randomly divided into two disjoint groups: 276 reference controls used to estimate normalization parameters and 512 pseudo-patients used for simulation. The simulated observations retained anatomical variability and cross-ROI covariance from the pseudo-patients’ baseline measurements. We simulated three progression subtypes, each affecting 15 of 96 FreeSurfer ROIs, comprising 5 shared and 10 subtype-specific regions. The resulting dataset contained 1,474 irregularly sampled longitudinal visits from 512 pseudo-patients, with known subtypes, stages, affected regions, and monotonic biomarker trajectories. Additional generation details are provided in Appendix A.2.

#### Evaluation metrics

We evaluated subtype recovery using the adjusted Rand index (ARI) and stage recovery using Stage *ρ* and Longitudinal Stage *ρ*. Stage *ρ* was defined as the Spearman correlation between true and inferred stages within each ground-truth subtype, averaged equally across subtypes. Longitudinal Stage *ρ* applied the same calculation to stage changes between adjacent visits from the same individual. To jointly evaluate the performance of subtyping and staging, we used joint subtype–stage concordance (JSSC) scores over within-individual visit pairs, rewarding pairs for which both visits were correctly subtyped and their inferred stages were correctly ordered. Detailed definitions of these metrics are provided in Appendix A.3.

### Comparison with existing methods

We compared TRACE with three cross-sectional clustering methods (K-Means, CHIMERA (Dong et al., 2015), and SmileGAN (Yang et al., 2021)) and four progression-aware methods (Surreal-GAN (Yang et al., 2022), SubLign (Chen et al., 2022), s– SuStaIn (Tandon et al., 2024)), and DPMoSt (Viani et al., 2025). All methods were evaluated under both cross-sectional and longitudinal settings where applicable. For each method, we performed ten runs with different initializations on the same semi-simulated dataset and report the mean and sample standard deviation.

As shown in Tab. 1, TRACE consistently achieved the strongest performance, substantially outper-forming the comparison methods in subtype recovery, disease staging, longitudinal stage-change estimation, and joint subtype–stage concordance. These results demonstrate the benefit of jointly modeling disease subtype and stage and show that TRACE effectively leverages longitudinal information while maintaining strong performance when only baseline observations are available.

**Table 1:**
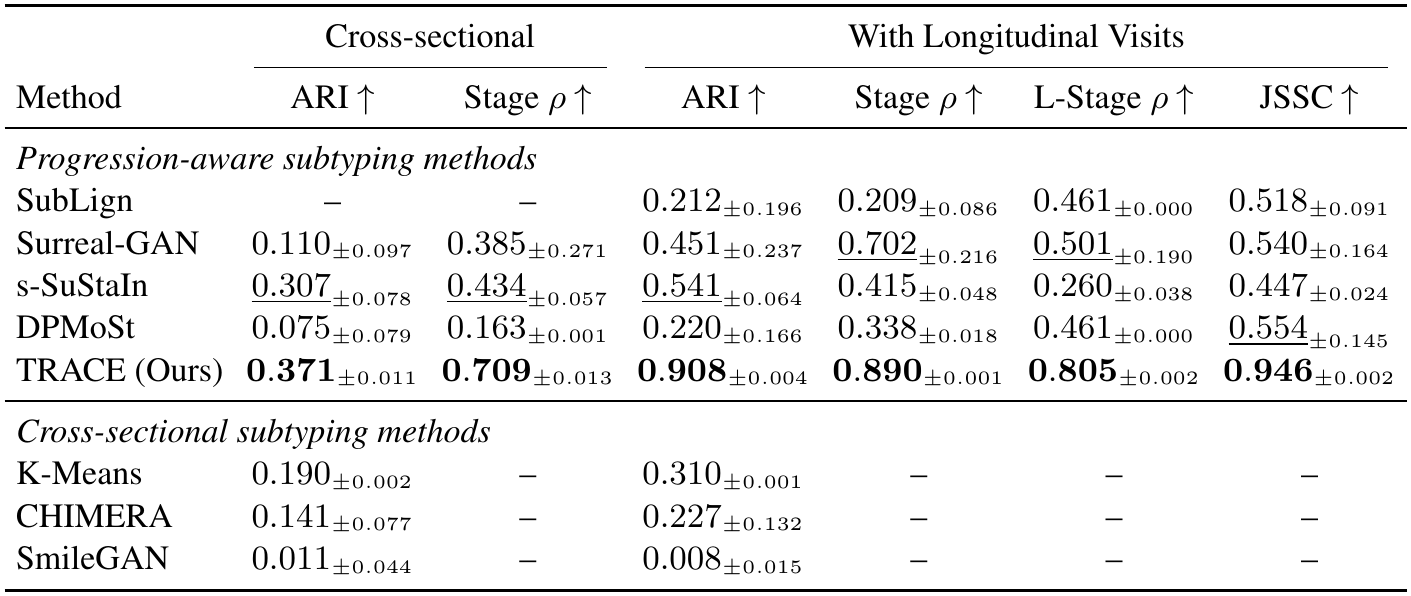
Subtype and stage recovery on the semi-simulated dataset under cross-sectional and longitudinal settings, using baseline visits only and all available visits, respectively. Longitudinal Stage *ρ* (L-Stage *ρ*) and JSSC are reported only for the longitudinal setting as they require longitudinal data. Methods without scalar stage estimates are evaluated using ARI only; SubLign is evaluated only in the longitudinal setting since it requires time-series inputs. Values are mean *±* sample standard deviation across ten random initializations. Best and second-best results are shown in bold and underlined, respectively.

| Method | Cross-sectional |  | With Longitudinal Visits |  |  |  |
|---|---|---|---|---|---|---|
|  | ARI ↑ | Stage ρ ↑ | ARI ↑ | Stage ρ ↑ | L-Stage ρ ↑ | JSSC ↑ |
| *Progression-aware subtyping methods* |  |  |  |  |  |  |
| SubLign | – | – | 0.212<sub>±0.196</sub> | 0.209<sub>±0.086</sub> | 0.461<sub>±0.000</sub> | 0.518<sub>±0.091</sub> |
| Surreal-GAN | 0.110<sub>±0.097</sub> | 0.385<sub>±0.271</sub> | 0.451<sub>±0.237</sub> | <u>0.702</u><sub>±0.216</sub> | <u>0.501</u><sub>±0.190</sub> | 0.540<sub>±0.164</sub> |
| s-SuStaIn | <u>0.307</u><sub>±0.078</sub> | <u>0.434</u><sub>±0.057</sub> | <u>0.541</u><sub>±0.064</sub> | 0.415<sub>±0.048</sub> | 0.260<sub>±0.038</sub> | 0.447<sub>±0.024</sub> |
| DPMoSt | 0.075<sub>±0.079</sub> | 0.163<sub>±0.001</sub> | 0.220<sub>±0.166</sub> | 0.338<sub>±0.018</sub> | 0.461<sub>±0.000</sub> | <u>0.554</u><sub>±0.145</sub> |
| TRACE (Ours) | **0.371**<sub>±0.011</sub> | **0.709**<sub>±0.013</sub> | **0.908**<sub>±0.004</sub> | **0.890**<sub>±0.001</sub> | **0.805**<sub>±0.002</sub> | **0.946**<sub>±0.002</sub> |
| *Cross-sectional subtyping methods* |  |  |  |  |  |  |
| K-Means | 0.190<sub>±0.002</sub> | – | 0.310<sub>±0.001</sub> | – | – | – |
| CHIMERA | 0.141<sub>±0.077</sub> | – | 0.227<sub>±0.132</sub> | – | – | – |
| SmileGAN | 0.011<sub>±0.044</sub> | – | 0.008<sub>±0.015</sub> | – | – | – |

#### Effectiveness of the change reconstruction loss

To determine an appropriate weight for the change reconstruction loss, we evaluated *λ*_chg_ ∈ {0, 10, 20, 30, 40, 50, 60} using ten different initializations of the same semi-simulated dataset. As shown in Fig. 2, performance improved substantially as the weight increased from 0 to 30, with weight 30 providing the best overall performance across most metrics. Relative to the model without the change reconstruction loss (*λ*_chg_ = 0), the selected weight improved all metrics in every run. These results demonstrate that the change reconstruction loss provides an effective longitudinal constraint and improves both the accuracy and robustness of disease-progression recovery.

**Figure 2:**
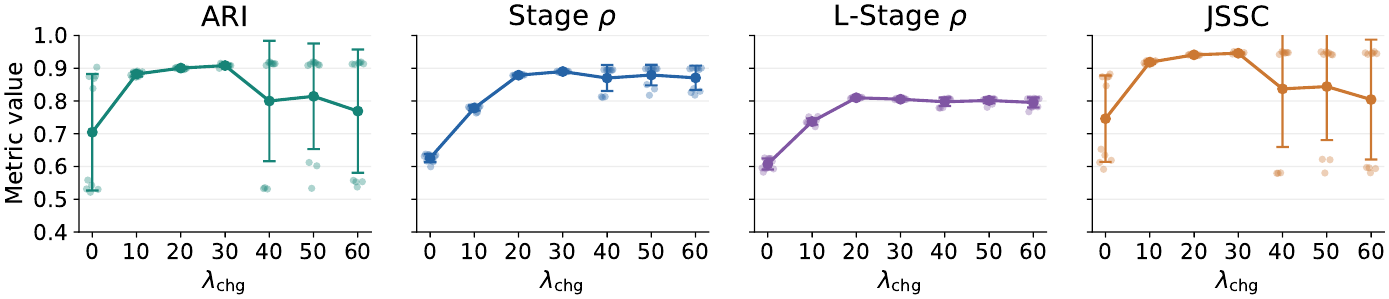
Sensitivity to the change-reconstruction weight on the semi-simulated dataset. Translucent points represent individual runs; solid points and error bars represent the mean and sample standard deviation across ten initializations.

We further visualized the fitted progression trajectories for one ROI affected across subtypes and one subtype-specific ROI for each subtype (Fig. 3). Without the change reconstruction loss, the estimated trajectories deviated considerably from the ground truth and exhibited spurious progression in ROIs which were unaffected. In contrast, using *λ*_chg_ = 30 yielded trajectories that more accurately recovered the timing, magnitude, and subtype specificity of the true progression patterns. This qualitative comparison is consistent with the quantitative results and further supports the effectiveness of the change reconstruction loss in recovering longitudinal disease progression.

**Figure 3:**
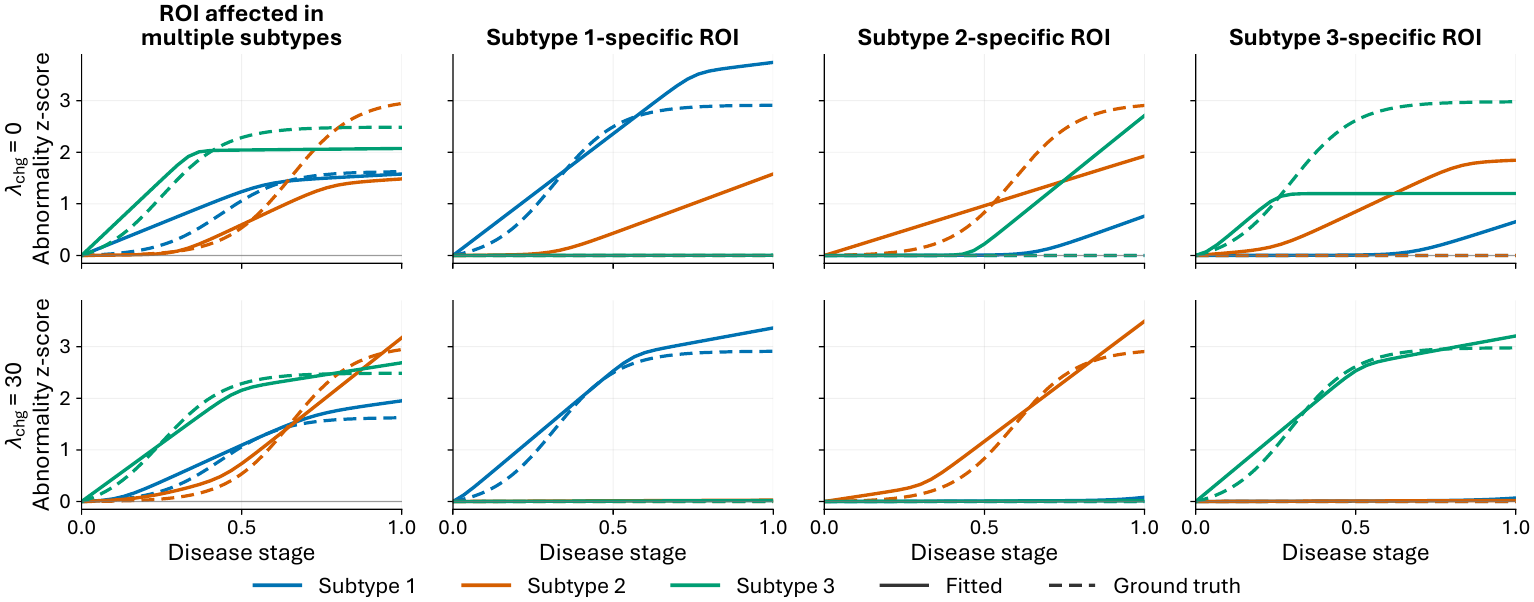
Fitted and ground-truth progression trajectories without the change reconstruction loss (top row, *λ*_chg_ = 0) and with the selected weight of 30 (bottom row). The columns show one ROI affected in multiple subtypes and one subtype-specific affected ROI for each subtype. Colors denote subtypes, solid lines represent fitted curves, and dashed lines represent ground truth.

### 3.3 Real-world Data Experiments

#### Data selection

We pooled individuals with diagnoses of CN, MCI, or dementia from ADNI and OASIS–3 who had complete measurements for all 96 MRI ROIs. A fixed reference group of 788 amyloid-negative baseline CN individuals (385 from ADNI and 403 from OASIS–3) was used to estimate age/sex adjustment and control standardization. All visits from reference individuals were excluded from model fitting and subtype discovery. The remaining discovery cohort comprised 2,732 participants and 9,721 visits.

#### Selecting the number of subtypes

We evaluate *K* = 2, …, 6 using five-fold subject-level cross-validation stratified by cohort and baseline diagnosis, with ten initializations for each *K*. For each initialization, models were trained on four folds and evaluated on the held-out fold. Log-likelihoods were summed across the five folds, normalized by the number of held-out visits, and then averaged across initializations. To assess stability, we computed the mean adjusted Rand index (ARI) across all pairs of initializations. We also summarized the subtype composition for each value of *K*.

The held-out log-likelihood improved steadily from *K* = 2 to *K* = 6, whereas across-initialization stability declined (Fig. 4(a),(b)). At *K* = 4, the mean log-likelihood was −133.53 per visit and the mean ARI was 0.659. Each of the four subtypes represented a substantial share of participants (18.2–40.4%; Fig. 4(c)). Considering predictive fit, stability, and the additional heterogeneity captured by the subtypes, we selected *K* = 4 for refitting on the full cohort.

**Figure 4:**
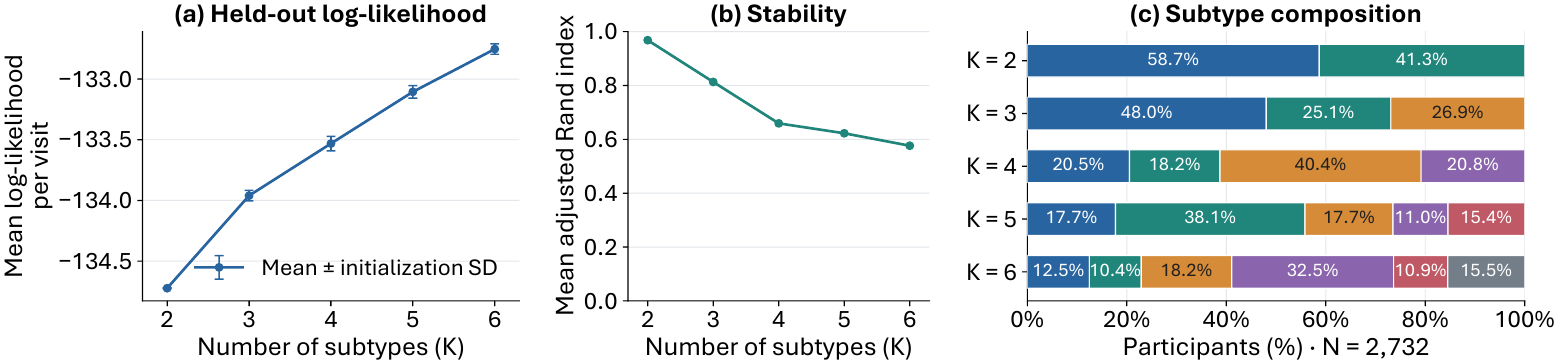
Real-world data model selection for the number of subtypes *K*. (a) Held-out marginal log-likelihood across ten initializations, with error bars indicating standard deviation. (b) Mean adjusted Rand index (ARI) across ten initializations. (c) Subtype percentages across out-of-fold assignments.

#### Refitting on the discovery set

We refit the model with *K* = 4 based on all 2,732 discovery participants using ten different initializations. We selected the run with the highest mean pairwise agreement in subtype assignments with the other runs. The fitted trajectories (Fig. 5) reveal distinct patterns of regional abnormality. Subtype 1 shows early medial temporal and subcortical abnormalities, followed by broader cortical involvement. Subtype 2 also shows early subcortical involvement, but its cortical involvement emerges later. Subtype 3 exhibits little visible abnormality until later inferred stages, whereas subtype 4 develops widespread cortical abnormalities by stage 0.4. These patterns broadly correspond to the clinical profiles (Tab. 2): subtype 1 has the lowest adjusted MMSE (25.72), highest CDR-SB (2.80) and FAQ (7.93), and fastest MMSE decline (−0.79 points/year); Subtype 3 has the mildest impairment and slowest decline; subtypes 2 and 4 have intermediate profiles, despite subtype 4’s earlier cortical involvement on the fitted stage scale.

**Figure 5:**
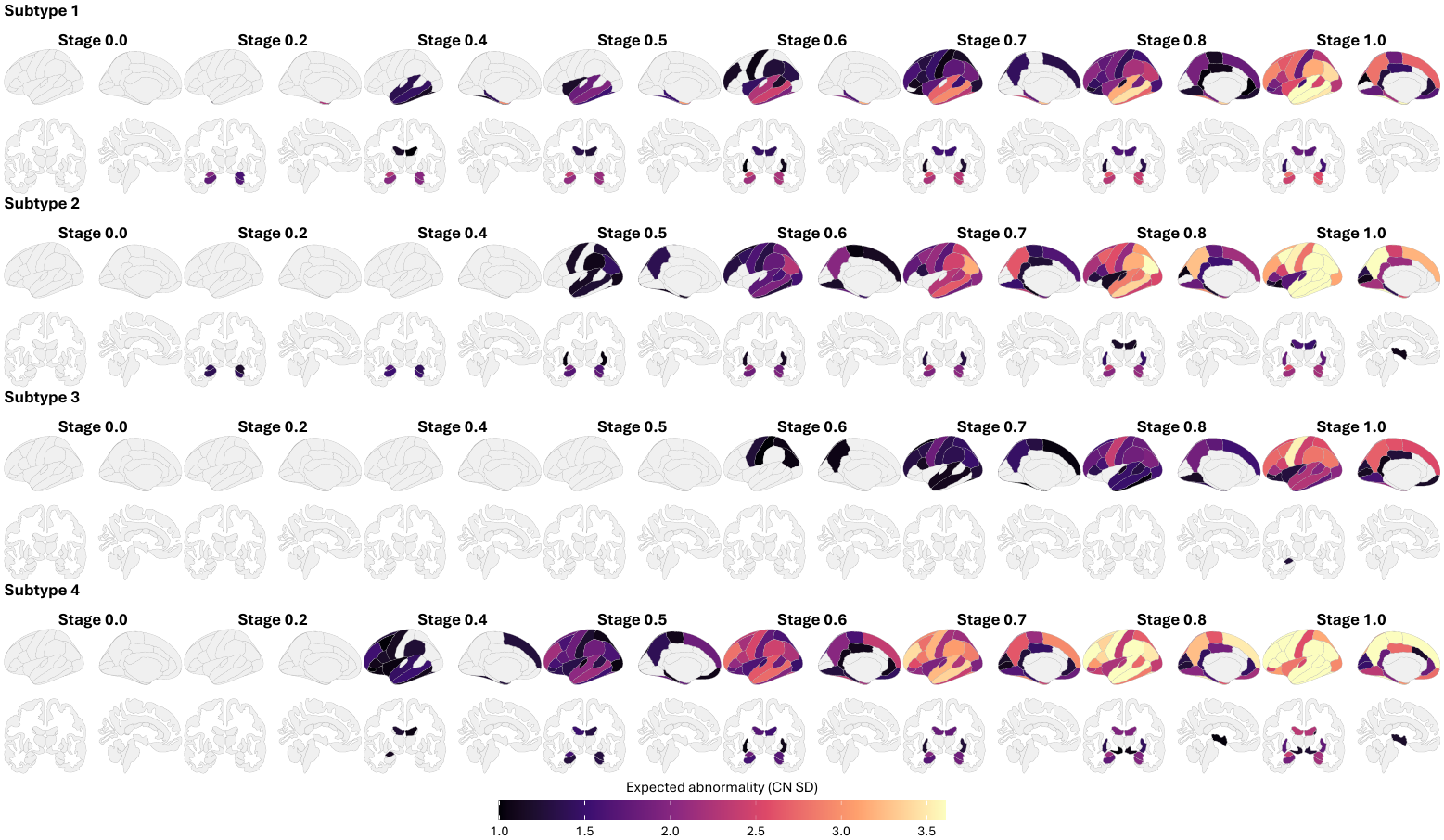
Subtype-specific progression trajectories on the full cohort with *K* = 4. Columns represent disease stage from 0.0 to 1.0. For each subtype, both cortical and subcortical views are shown. Colors indicate expected abnormality in units of reference-control standard deviations.

**Table 2:** Clinical associations of the four MRI-derived progression subtypes. Baseline cognitive scores were analyzed using regression models that included subtype, inferred stage, age, sex, education, and cohort. Annual changes in cognitive scores were estimated from repeated observations using baseline subtype and stage, with adjustment for age, sex, education, and cohort. Parentheses indicate 95% confidence intervals in the *Baseline clinical associations* and *Longitudinal clinical associations*. The *p*-value column reports statistical tests of overall subtype differences in baseline clinical scores or annual rates of change.

| Measure | Subtype 1 | Subtype 2 | Subtype 3 | Subtype 4 | *p*-value |
|---|---|---|---|---|---|
| *Cohort and model* |  |  |  |  |  |
| Participants, *n* | 453 | 660 | 1073 | 546 | — |
| Baseline CN, *n* (%) | 43 (9.5%) | 189 (28.6%) | 598 (55.7%) | 158 (28.9%) | — |
| Baseline MCI, *n* (%) | 194 (42.8%) | 269 (40.8%) | 382 (35.6%) | 275 (50.4%) | — |
| Baseline Dementia, *n* (%) | 216 (47.7%) | 202 (30.6%) | 93 (8.7%) | 113 (20.7%) | — |
| *Baseline clinical associations* |  |  |  |  |  |
| MMSE | 25.72 (25.46, 25.98) | 26.71 (26.52, 26.89) | 28.36 (28.24, 28.47) | 27.13 (26.91, 27.35) | < 0.001 |
| CDR-SB | 2.80 (2.61, 2.99) | 1.88 (1.75, 2.00) | 0.78 (0.71, 0.85) | 1.57 (1.43, 1.71) | < 0.001 |
| FAQ | 7.93 (7.25, 8.62) | 4.80 (4.33, 5.27) | 1.71 (1.46, 1.96) | 4.30 (3.77, 4.83) | < 0.001 |
| *Longitudinal clinical associations* |  |  |  |  |  |
| MMSE change per year | -0.79 (-0.87, -0.70) | -0.41 (-0.45, -0.37) | -0.13 (-0.16, -0.11) | -0.35 (-0.41, -0.28) | < 0.001 |
| CDR-SB change per year | 0.64 (0.58, 0.69) | 0.34 (0.31, 0.36) | 0.10 (0.08, 0.11) | 0.25 (0.22, 0.29) | < 0.001 |

## 4 Conclusion

We presented TRACE, a probabilistic model that disentangles disease subtypes from stages using cross-sectional MRI observations and short-term longitudinal changes when available. Its monotonic neural network based trajectories flexibly capture nonlinear biomarker progression, while end-to-end optimization enables computationally efficient and scalable estimation. On semi-simulated data, TRACE improved subtype and stage recovery over exisiting methods in both cross-sectional and longitudinal settings, and incorporating observed longitudinal changes further improved recovery of the underlying trajectories. Applied to real-world datasets, TRACE identified four anatomically distinct progression patterns associated with differences in clinical impairment.

## AI use statement

In this work, we used generative AI tools to polish the writing, assist with method implementation, the presentation and consistency checking of equations, and the implementation of semi-simulated data generation. We did not use generative AI to preprocess the source cohort data, design the methodology or experiments, draft sectionals of the paper or generate plots. Other tasks requiring disclosure were not applicable. We reviewed and verified all AI-assisted material and take full responsibility for the final content of this work, including its text, claims, code, and plots.

## Ethics statement

This study analyzes publicly available neuroimaging and clinical data from ADNI and OASIS–3. It does not recruit participants or intervene in their care. We report aggregate findings and do not include participant-level images or clinical records in this manuscript. Access to the underlying cohort data is governed by the respective data providers.

## Reproducibility statement

The observation model, subtype and stage inference, trajectory parameterization, and longitudinal objective are specified in Sec. 2, with network architecture and optimization settings in Sec. 2.6. The Experiments and Results section describes cohort selection, MRI preprocessing, baselines, evaluation settings, and model selection. Appendix A.2 gives the semi-simulated data-generation procedure, and Appendix A.3 defines the recovery metrics. We plan to provide source code as well as the semi-simulated dataset for reproducing the semi-simulated results. Reproducing the real-data analysis requires access to ADNI and OASIS–3 under their respective data-use terms.

## A Appendix: Additional experimental details

### A.1 Details of Real Dataset

**Table S1:** Characteristics of datasets. Age, sex, education, and diagnosis are summarized at baseline. Visits and follow-up are median [IQR]; follow-up includes participants with at least two visits. Other values are *n* (%) or mean *±* standard deviation.

| Characteristic | ADNI | OASIS–3 | Combined |
|---|---|---|---|
| Participants, *n* | 2,363 | 1,157 | 3,520 |
| Imaging visits, *n* | 9,656 | 2,113 | 11,769 |
| Age, years | 73.24 ± 7.32 | 70.45 ± 8.84 | 72.33 ± 7.96 |
| Female sex, *n* (%) | 1,120 (47.4) | 644 (55.7) | 1,764 (50.1) |
| Education, years | 16.04 ± 2.74 | 15.85 ± 2.67 | 15.98 ± 2.72 |
| Baseline CN, *n* (%) | 871 (36.9) | 905 (78.2) | 1,776 (50.5) |
| Baseline MCI, *n* (%) | 1,082 (45.8) | 38 (3.3) | 1,120 (31.8) |
| Baseline dementia, *n* (%) | 410 (17.4) | 214 (18.5) | 624 (17.7) |
| Visits per participant | 4 [2–5] | 1 [1–2] | 3 [1–5] |
| Participants with ≥ 2 visits, *n* (%) | 1,931 (81.7) | 522 (45.1) | 2,453 (69.7) |
| Follow-up, years | 2.61 [1.92–5.05] | 4.40 [2.78–7.18] | 3.01 [1.97–6.01] |

### A.2 Details of semi-simulated dataset generation

#### Normalization

To retain realistic anatomical variation on a common scale, we regressed each raw ROI measurement on centered age and sex in the reference controls. We divided each pseudopatient’s residual by the corresponding reference residual standard deviation, aligned its direction so that positive values indicated greater abnormality, and clipped it to [ −8, 8]. Simulation therefore took place in the normalized *z*-score domain.

#### Subtype and affected-ROI assignment

To establish known heterogeneity, we assigned pseudopatients to three subtypes in proportions 4 : 3 : 2, yielding 227, 171, and 114 participants. Each subtype affected 15 ROIs: five were shared across all three subtypes, and ten were unique to that subtype. The shared ROIs were randomly sampled once, and the subtype-specific sets were mutually disjoint. Assignments and affected-ROI sets remained fixed across visits.

#### Progression trajectories

To vary the timing and magnitude of regional abnormalities, we randomly sampled a separate logistic trajectory for every affected subtype–ROI pair. Its endpoint amplitude was uniform on [1.5, 3.0] reference standard deviations, its transition center was uniform on [0.15, 0.85], and its width was 0.10. Each curve was rescaled to equal zero at stage 0 and its sampled amplitude at stage 1. Even for shared ROIs, trajectories were sampled independently across subtypes. Unaffected subtype–ROI pairs had no disease-related signal.

#### Visit schedules and disease stages

To emulate irregular, short-term follow-up, we sampled the visit count and initial stage for participant *i* as

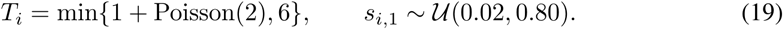

Inter-visit intervals Δ*t*_*i,t*_ (in years) and progression multipliers *v*_*i,t*_ were sampled independently from log-normal distributions, each with mean one and log-scale standard deviation 0.35. For each subsequent visit, the proposed stage increment was

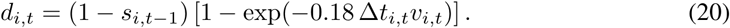

We capped the increment at 0.20, imposed a numerical minimum of 10^−5^, and truncated the updated stage at 1. This produced monotone progression that slows as the stage approaches 1.

#### Simulated observations

To obtain each visit’s ROI measurement, we combined the baseline measurement, the disease signal for the assigned subtype *k*_*i*_ at the visit’s stage, and session noise:

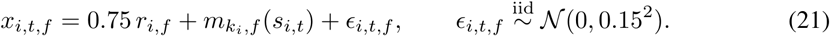

### A.3 Details of evaluation metrics

#### Subtype recovery

We computed the adjusted Rand index (ARI) between the true and inferred session-level partitions. ARI is invariant to subtype-label permutations, adjusts for chance agreement, and equals one for perfect recovery.

#### Stage recovery

Stage *ρ* was computed as the Spearman correlation between true and inferred stages within each true subtype, averaged with equal weight across subtypes. Sessions were grouped by their true subtype regardless of their inferred assignment.

For Longitudinal Stage *ρ*, we ordered visits within each participant and calculated true and inferred stage changes between adjacent visits. We then computed the Spearman correlation between these changes within each true subtype and averaged the correlations with equal weight across subtypes. This metric evaluates stage changes and was omitted for cross-sectional data.

#### Joint subtype–stage concordance

To jointly evaluate the subtype assignment and stage ordering, we first aligned inferred subtype labels to the true labels, denoting the aligned assignments by 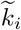 Let *P*_*k*_ contain all within-participant visit pairs of true subtype *k* with unequal true stages, oriented so that *s*_*i*_ *< s*_*j*_. Each pair received the score

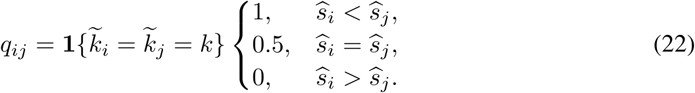

Thus, a pair received credit only when both visits were correctly subtyped, with full credit for correct stage ordering and half credit for an inferred stage tie. JSSC was obtained by averaging pair scores within each subtype and then equally across subtypes:

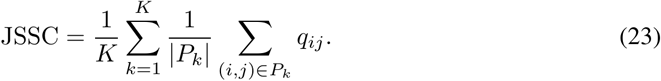

JSSC ranges from zero to one, with one indicating correct subtype assignments and stage ordering for all evaluated pairs.

### A.4 Implementation of comparison methods

All methods were fit to the same semi-simulated dataset with three subtypes and 96 ROIs. The longitudinal setting contained 1,474 visits from 512 participants. The cross-sectional setting retained only each participant’s baseline visit. Inputs were age- and sex-adjusted, control-standardized ROI measurement. True subtypes and stages were used only for evaluation. Cross-sectional clustering methods without a scalar stage estimate were evaluated for subtype recovery alone.

#### K-Means

We applied K-Means with three clusters directly to the ROI vectors. In the longitudinal setting, visits were clustered independently. K-Means does not estimate disease stages.

#### CHIMERA

We used the authors’ affine-transformation model with three subtype components. Each run used three optimization starts, and we retained the start with the highest mean agreement with the other starts. CHIMERA does not estimate disease stages.

#### SmileGAN

We used the authors’ released implementation and set the cluster number to 3, the cluster-loss weight to 9, the change-loss weight to 5, and batch size 256. When the released stopping criterion was unmet, we evaluated the final checkpoint. Each visit was treated as an independent observation. SmileGAN does not estimate disease stages.

#### Surreal-GAN

We used the authors’ released implementation with four training repetitions per initialization. It produced three continuous disease-related R-indices per observation. We assigned the subtype by the largest R-index and used that maximum as a prespecified severity indicator for stage-related evaluations.

#### SubLign

We supplied each participant’s ordered, padded visit sequence and elapsed follow-up times to the released longitudinal model. Its bounded decoder required feature-wise min–max scaling of the common ROI measurements to [0, 1]. We trained for 200 epochs, inferred one subtype per participant, and formed the stage ranking from elapsed time plus the inferred participant-specific time shift. We evaluated SubLign only in the longitudinal setting and did not perform a baseline-only experiment.

#### s–SuStaIn

For the 96-ROI experiments, we used s-SuStaIn, a scalable extension of SuStaIn that groups biomarkers to reduce the computational cost of event-order inference (Tandon et al., 2024). We fitted three subtypes with five biomarker clusters. Optimization used ten starts and at most 50 iterations. We estimated stage as the maximum-posterior event position divided by the number of clusters.

#### DPMoSt

DPMoSt was evaluated using the authors’ implementation with support for three subtype components. Biomarkers were rescaled to [0, 3] and elapsed follow-up times to [0, 20]. We retained the released initialization and optimization budget: 1,000 initialization steps per biomarker, 30 outer iterations with 30 updates per parameter group, and 50 final E steps. Native subtype assignments and estimated times were evaluated.

## B Appendix: Additional Experiments and Results

### B.1 Generalization of cross-sectional fitting

In order to further evaluate models’ ability to recovery progression heterogeneity on cross-sectional data, we fit TRACE, s–SuStaIn, Surreal-GAN and DPMoSt using baseline scans, then apply the fitted models to all follow-up scans independently. We evaluate this design on the both semi-simulated dataset and real discovery dataset. All methods use *K* = 3 on semi-simulated data and use *K* = 4 on real data.

#### Semi-simulated data

We evaluate 962 follow-up visits from 454 participants. We calculated ARI, stage *ρ*, Longitudinal stage *ρ* and JSSC. We also report the fraction of participants whose every follow-up subtype assignment matches their baseline assignment (Tab. S2). TRACE has the strongest mean stage recovery, JSSC and consistent subtyping results. s-SuStaIn has a slightly higher mean follow-up ARI but substantially more variation across repeated runs.

#### Real data

The discovery dataset contains 2,732 participants, of whom 1,971 have 6,989 follow-up scans. Ground-truth subtypes and stages are unavailable, so we report subtype consistency instead, including mean within-participant follow-up match and the fraction whose every follow-up matches baseline. Table S3 shows TRACE has the highest consistency.

**Table S2:** Follow-up visit inference after cross-sectional fitting on the semi-simulated dataset. All follow-ups match is the fraction of participants whose every follow-up subtype assignment matches baseline. Values are mean *±* sample standard deviation over ten initializations.

| Method | Follow-up ARI ↑ | Follow-up Stage ρ ↑ | L-Stage ρ ↑ | JSSC ↑ | All follow-ups match ↑ |
|---|---|---|---|---|---|
| TRACE (Ours) | 0.545 ± 0.008 | **0.580 ± 0.033** | **0.642 ± 0.012** | **0.549 ± 0.030** | **0.889 ± 0.009** |
| s-SuStaIn | **0.564 ± 0.104** | 0.279 ± 0.053 | 0.201 ± 0.059 | 0.366 ± 0.032 | 0.826 ± 0.021 |
| Surreal-GAN | 0.192 ± 0.164 | 0.370 ± 0.218 | 0.168 ± 0.224 | 0.294 ± 0.089 | 0.856 ± 0.059 |
| DPMoSt | 0.381 ± 0.153 | 0.371 ± 0.002 | 0.456 ± 0.000 | 0.299 ± 0.168 | 0.433 ± 0.136 |

**Table S3:** Subtype consistency across follow-up visits in the real discovery dataset. Mean follow-up match averages each participant’s fraction of follow-up scans assigned to their baseline subtype. All follow-ups match is the fraction of participants whose every follow-up assignment matches baseline. Values are mean *±* sample standard deviation across ten initializations.

| Method | Mean follow-up match ↑ | All follow-ups match ↑ |
|---|---|---|
| TRACE (Ours) | **0.748 ± 0.014** | **0.572 ± 0.017** |
| s-SuStaIn | 0.726 ± 0.011 | 0.549 ± 0.018 |
| Surreal-GAN | 0.685 ± 0.049 | 0.494 ± 0.062 |
| DPMoSt | 0.235 ± 0.140 | 0.117 ± 0.085 |

**Table S4:** Results of different trajectory function on the semi-simulated longitudinal dataset. Values are mean sample standard deviation over ten random initializations. All models use a change reconstruction loss weight of 30.

| Trajectory family | ARI ↑ | Stage *ρ* ↑ | L-Stage *ρ* ↑ | JSSC ↑ | Trajectory RMSE ↓ |
|---|---|---|---|---|---|
| SMM | **0.908 ± 0.004** | **0.890 ± 0.001** | **0.805 ± 0.002** | **0.946 ± 0.002** | **0.081 ± 0.002** |
| Piecewise-linear | 0.597 ± 0.197 | 0.808 ± 0.041 | 0.743 ± 0.038 | 0.776 ± 0.100 | 0.250 ± 0.063 |
| I-spline | 0.496 ± 0.128 | 0.815 ± 0.019 | 0.729 ± 0.023 | 0.649 ± 0.147 | 0.336 ± 0.103 |

### B.2 Alternative Parametric Trajectory Functions

We compared the SMM trajectory network with two lower-dimensional monotonic families in the same subtype–stage likelihood and change-reconstruction framework. For subtype *k* and ROI *f*, both alternatives take the form

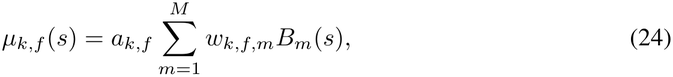

where *M* = 7 and *B*_*m*_ is a basis function. The learned weights *w*_*k,f,m*_ are obtained by a softmax transformation, making them nonnegative and summing to one. They allocate the total trajectory increase across the basis functions. Their cumulative contributions ensure that abnormality cannot decrease with stage. The scale *a*_*k,f*_ is the learned endpoint for this subtype–ROI pair. Thus, each trajectory begins at zero and reaches its learned endpoint at stage one.

#### Piecewise-linear trajectories

We divide the stage range into *M* = 7 equal segments and assign one cumulative ramp basis function to each segment. A ramp is zero before its segment, rises linearly across that segment, and remains one afterward. Summing the weighted ramps gives a continuous piecewise-linear trajectory whose slope can change at segment boundaries.

#### I-spline trajectories

We instead use normalized cubic I-spline basis functions, formed by integrating nonnegative quadratic B-spline functions over stage. Each I-spline increases smoothly from zero to one, with overlapping basis functions allowing gradual changes in slope. Their nonnegative weighted sum is therefore smooth and monotonic, with the same zero and learned endpoint anchors as the piecewise-linear trajectory.

#### Results

SMM showed the best overall subtype and stage recovery, the lowest trajectory error, and the most consistent performance across initializations (Tab. S4). In Fig. 6, SMM followed the simulated shared and subtype-specific trajectories more closely. The alternatives showed larger departures in curve shape or endpoint.

**Figure 6:**
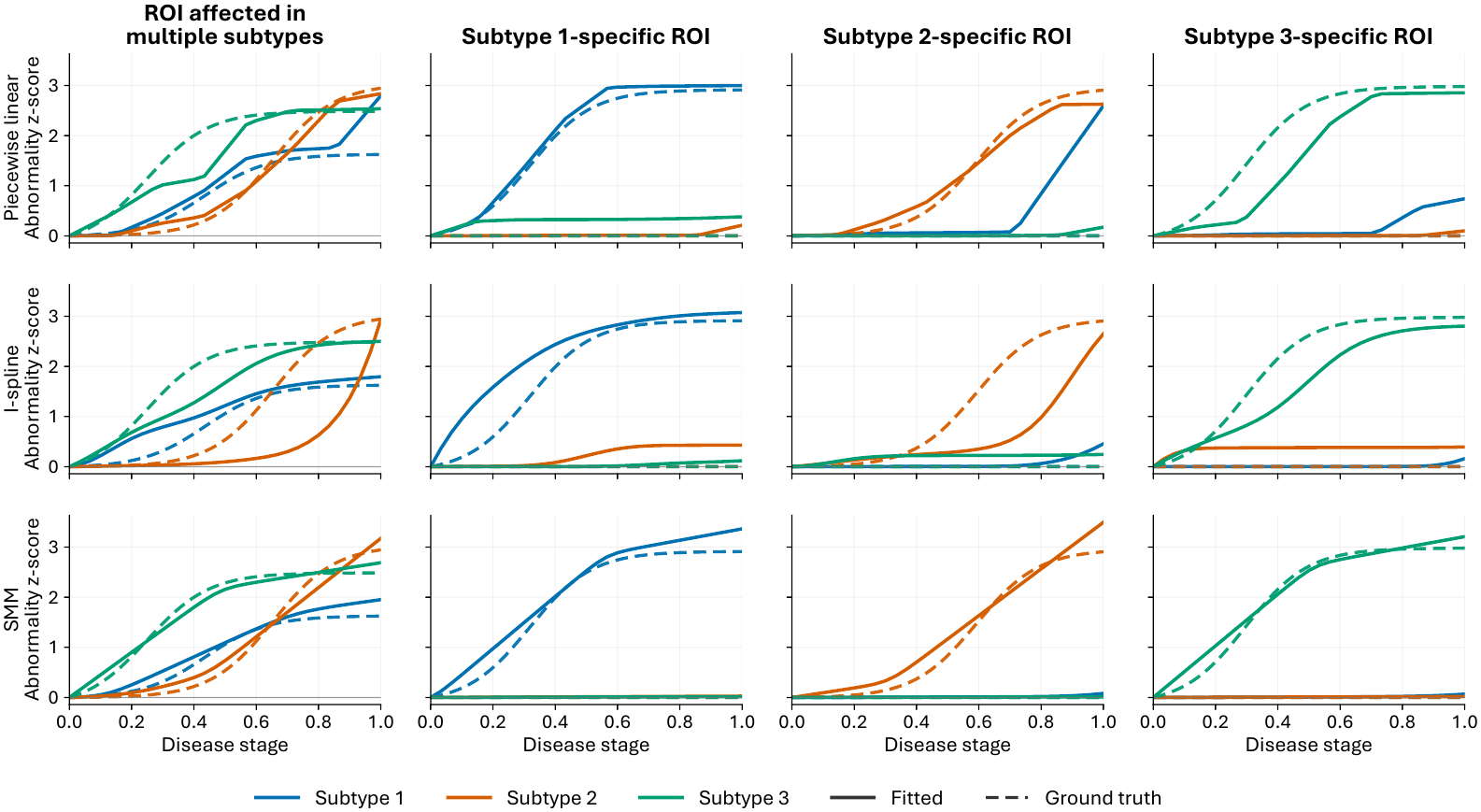
Fitted and ground-truth progression curves of piecewise-linear (top row), I-spline (middle row) and SMM (bottom row). Columns show one ROI affected in multiple subtypes and one subtype-specific ROI for each subtype. Colors denote subtypes, solid lines represent fitted curves, and dashed lines represent ground truth.

### B.3 Visualization of ROI Trajectories on Real Data

We examined the full-discovery *K* = 4 refit on specific ROIs. ROIs were ranked by the mean absolute difference across the fitted subtype curves, averaged over the disease-stage grid. The ten highest-ranked ROIs (Fig. 7) comprise bilateral entorhinal, temporal-pole, and middle-temporal cortical regions, as well as bilateral amygdala and hippocampus. Subtype 1 shows the earliest and largest abnormalities in several medial temporal ROIs, whereas subtype 3 has lower, later-rising trajectories. Subtypes 2 and 4 generally lie between these patterns but differ in the timing and magnitude of regional change.

**Figure 7:**
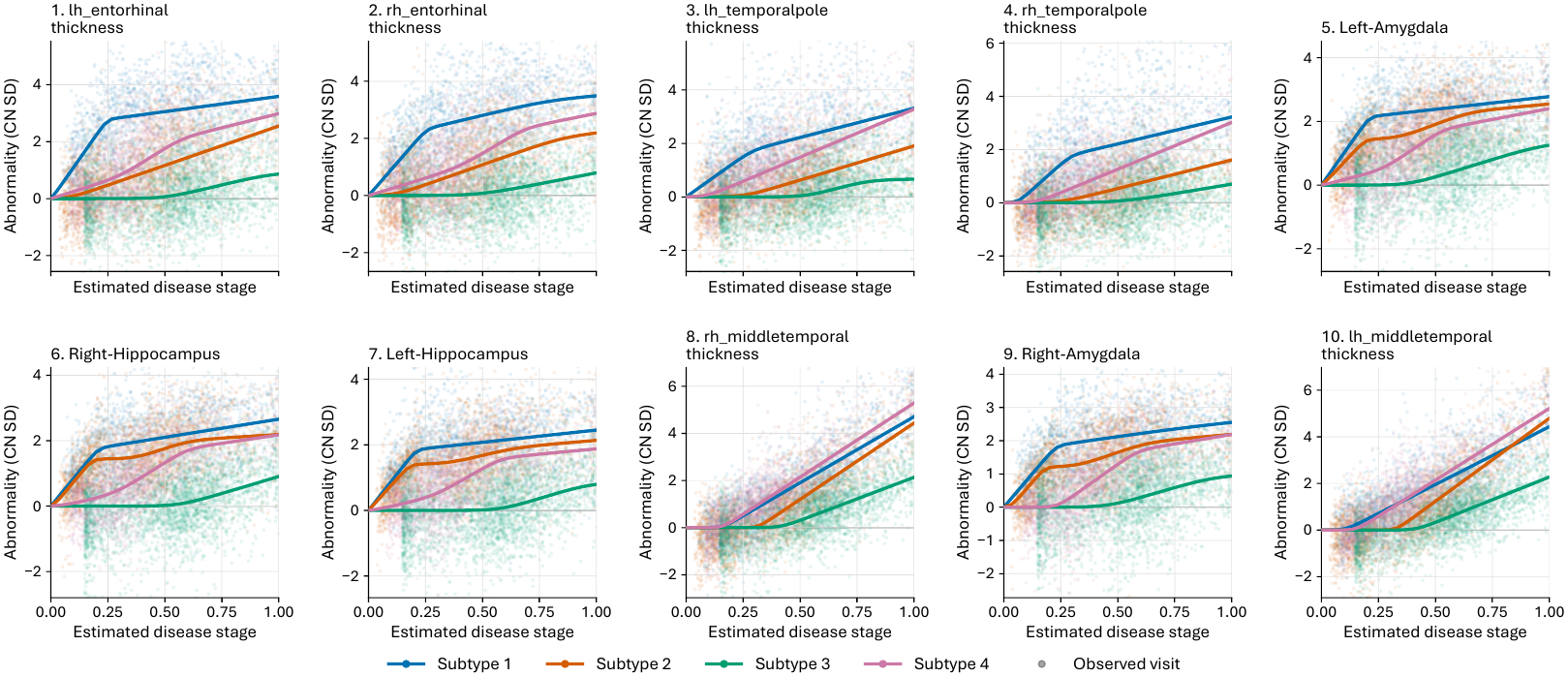
Ten ROIs with the greatest stage-averaged mean pairwise separation among fitted curves in the *K* = 4 real-data refit. Solid lines show subtype-specific expected abnormalities in reference-control standard-deviation units. Points show observed visits, colored by inferred subtype and positioned at inferred disease stage.

### B.4 Comparison with SuStaIn

**Table S5:** Performance on pySuStaIn-generated data (mean±standard deviation across three simulation seeds). Higher ARI and stage correlation are better; lower runtime is better.

| Method | Case ARI | Stage ρ (all) | Stage ρ (cases) | Runtime (s) |
|---|---|---|---|---|
| TRACE | 0.3399 ± 0.1208 | 0.9541 ± 0.0016 | 0.9423 ± 0.0030 | 18.8 ± 5.0 |
| z-score SuStaIn | 0.3473 ± 0.1775 | 0.9519 ± 0.0039 | 0.9373 ± 0.0055 | 900.9 ± 23.9 |

Our main 96-ROI experiments use s-SuStaIn, a scalable extension of SuStaIn (Tandon et al., 2024). To additionally compare with the original SuStaIn algorithm (Young et al., 2018), we generated three cross-sectional datasets with five biomarkers using the simulation procedure provided in pySuStaIn (Aksman et al., 2021) (https://github.com/ucl-pond/pySuStaIn/blob/master/sim/simfuncs.py), with a different random seed for each dataset.

Each dataset contained 800 subjects, 5 biomarkers, and 3 disease subtypes; 25% of subjects were stage-zero controls. We then fitted TRACE and z-score SuStaIn to the same observations, specifying three subtypes for both methods. Subtype recovery was measured by adjusted Rand index (ARI) among cases, and stage recovery by Spearman correlation with the simulated stage. We also recorded runtime for each method on CPU. Table S5 reports means and sample standard deviations across the three datasets. TRACE achieved similar subtype and stage recovery to z-score SuStaIn on these datasets, with substantially shorter runtime.

## References

Emma Ahlqvist, Petter Storm, Annemari Käräjämäki, Mats Martinell, Mozhgan Dorkhan, Annelie Carlsson, Petter Vikman, Rashmi B Prasad, Dina Mansour Aly, Peter Almgren, et al. Novel sub-groups of adult-onset diabetes and their association with outcomes: a data-driven cluster analysis of six variables. The lancet Diabetes & endocrinology, 6(5):361–369, 2018. 1

Leon M Aksman, Peter A Wijeratne, Neil P Oxtoby, Arman Eshaghi, Cameron Shand, Andre Altmann, Daniel C Alexander, and Alexandra L Young. pysustain: a python implementation of the subtype and stage inference algorithm. SoftwareX, 16:100811, 2021. 17

Randall J. Bateman, Chengjie Xiong, Tammie L. S. Benzinger, et al. Clinical and biomarker changes in dominantly inherited Alzheimer’s disease. New England Journal of Medicine, 367(9):795–804, 2012. 1

Irene Y. Chen, Rahul G. Krishnan, and David Sontag. Clustering interval-censored time-series for disease phenotyping. In Proceedings of the AAAI Conference on Artificial Intelligence, volume 36, pp. 6211–6221, 2022. 2, 7

Sterre de Jonge, Elisabeth J Vinke, Meike W Vernooij, Daniel C Alexander, Alexandra L Young, and Esther E Bron. Disease progression and subtype modeling for combined discrete and continuous input data. In 2026 IEEE 23rd International Symposium on Biomedical Imaging (ISBI), pp. 1–4, 2026. 2

Rahul S. Desikan, Florent Ségonne, Bruce Fischl, Brian T. Quinn, Bradford C. Dickerson, Deborah Blacker, Randy L. Buckner, Anders M. Dale, R. Paul Maguire, Bradley T. Hyman, Marilyn S. Albert, and Ronald J. Killiany. An automated labeling system for subdividing the human cerebral cortex on MRI scans into gyral based regions of interest. NeuroImage, 31(3):968–980, 2006. 6

Aoyan Dong, Nicolas Honnorat, Bilwaj Gaonkar, and Christos Davatzikos. Chimera: Clustering of heterogeneous disease effects via distribution matching of imaging patterns. IEEE transactions on medical imaging, 35(2):612–621, 2015. 2, 7

Michael C Donohue, Hélène Jacqmin-Gadda, Mélanie Le Goff, Ronald G Thomas, Rema Raman, Anthony C Gamst, Laurel A Beckett, Clifford R Jack Jr, Michael W Weiner, Jean-François Dartigues, et al. Estimating long-term multivariate progression from short-term data. Alzheimer’s & Dementia, 10:S400–S410, 2014. 1

Bruce Fischl. FreeSurfer. NeuroImage, 62(2):774–781, 2012. 6

Bruce Fischl, David H. Salat, Evelina Busa, Marilyn Albert, Megan Dieterich, Christian Haselgrove, Andre van der Kouwe, Ron Killiany, David Kennedy, Shuna Klaveness, Albert Montillo, Nikos Makris, Bruce Rosen, and Anders M. Dale. Whole brain segmentation: automated labeling of neuroanatomical structures in the human brain. Neuron, 33(3):341–355, 2002. 6

Christian Igel. Smooth min-max monotonic networks. In Proceedings of the 41st International Conference on Machine Learning, 2024. 2, 4

Sohyun Kang, Sung-Woo Kim, Joon-Kyung Seong, Alzheimer’s Disease Neuroimaging Initiative, et al. Disentangling brain atrophy heterogeneity in alzheimer’s disease: A deep self-supervised approach with interpretable latent space. NeuroImage, 297:120737, 2024. 2

Pamela J. LaMontagne, Tammie L. S. Benzinger, John C. Morris, et al. OASIS-3: Longitudinal neuroimaging, clinical, and cognitive dataset for normal aging and Alzheimer disease. medRxiv, 2019. 6

Ilya Loshchilov and Frank Hutter. Decoupled weight decay regularization. In International Conference on Learning Representations, 2019. 6

Susanne G. Mueller, Michael W. Weiner, Leon J. Thal, Ronald C. Petersen, Clifford R. Jack, William Jagust, John Q. Trojanowski, Arthur W. Toga, and Laurel Beckett. Ways toward an early diagnosis in Alzheimer’s disease: The Alzheimer’s Disease Neuroimaging Initiative (ADNI). Alzheimer’s & Dementia, 1(1):55–66, 2005. 6

Raghav Tandon, James J Lah, and Cassie S Mitchell. s-sustain: Scaling subtype and stage inference via simultaneous clustering of subjects and biomarkers. In Proceedings of the fifth Conference on Health, Inference, and Learning, volume 248, pp. 461–476, 2024. 2, 7, 15, 17

Erdem Varol, Aristeidis Sotiras, Christos Davatzikos, Alzheimer’s Disease Neuroimaging Initiative, et al. Hydra: Revealing heterogeneity of imaging and genetic patterns through a multiple max-margin discriminative analysis framework. Neuroimage, 145:346–364, 2017. 2

Alessandro Viani, Anna Custo, Emile d’Angremont, Valentina Garibotto, Giovanni B. Frisoni, Boris A. Gutman, and Marco Lorenzi. Disease progression modeling and stratification for detecting sub-trajectories in the natural history of pathologies: Application to alzheimer’s disease trajectory modeling. Imaging Neuroscience, 3:IMAG.a.954, 11 2025. 2, 7

Zhijian Yang, Ilya M Nasrallah, Haochang Shou, Junhao Wen, Jimit Doshi, Mohamad Habes, Guray Erus, Ahmed Abdulkadir, Susan M Resnick, Marilyn S Albert, et al. A deep learning framework identifies dimensional representations of alzheimer’s disease from brain structure. Nature communications, 12(1):7065, 2021. 2, 7

Zhijian Yang, Junhao Wen, and Christos Davatzikos. Surreal-GAN: Semi-supervised representation learning via GAN for uncovering heterogeneous disease-related imaging patterns. In International Conference on Learning Representations, 2022. 2, 7

Zhijian Yang, Junhao Wen, Guray Erus, Sindhuja T Govindarajan, Randa Melhem, Elizabeth Mamourian, Yuhan Cui, Dhivya Srinivasan, Ahmed Abdulkadir, Paraskevi Parmpi, et al. Brain aging patterns in a large and diverse cohort of 49,482 individuals. Nature medicine, 30(10):3015–3026, 2024. 2, 4

Alexandra L. Young, Razvan V. Marinescu, Neil P. Oxtoby, et al. Uncovering the heterogeneity and temporal complexity of neurodegenerative diseases with subtype and stage inference. Nature Communications, 9(1):4273, 2018. 2, 4, 17

Alexandra L. Young, Jacob W. Vogel, Leon M. Aksman, Peter A. Wijeratne, Arman Eshaghi, Neil P. Oxtoby, Steven C. R. Williams, and Daniel C. Alexander. Ordinal SuStaIn: Subtype and stage inference for clinical scores, visual ratings, and other ordinal data. Frontiers in Artificial Intelligence, 4:613261, 2021. 2

Alexandra L Young, Neil P Oxtoby, Sara Garbarino, Nick C Fox, Frederik Barkhof, Jonathan M Schott, and Daniel C Alexander. Data-driven modelling of neurodegenerative disease progression: thinking outside the black box. Nature Reviews Neuroscience, 25(2):111–130, 2024. 1

